# The phospho-docking protein 14-3-3 regulates microtubule-associated proteins in oocytes including the chromosomal passenger Borealin

**DOI:** 10.1101/2021.12.16.472885

**Authors:** Charlotte Repton, C. Fiona Cullen, Mariana F. A. Costa, Christos Spanos, Juri Rappsilber, Hiroyuki Ohkura

## Abstract

Global regulation of spindle-associated proteins is crucial in oocytes due to the absence of centrosomes and their very large cytoplasmic volume, but little is known about how this is achieved beyond involvement of the Ran-importin pathway. We previously uncovered a novel regulatory mechanism in *Drosophila* oocytes, in which the phospho-docking protein 14-3-3 suppresses microtubule binding of Kinesin-14/Ncd away from chromosomes. Here we report systematic identification of microtubule-associated proteins regulated by 14-3-3 from *Drosophila* oocytes. Proteins from ovary extract were co-sedimented with microtubules in the presence or absence of a 14-3-3 inhibitor. Through quantitative mass-spectrometry, we identified proteins or complexes whose ability to binding microtubules is suppressed by 14-3-3, including the chromosomal passenger complex (CPC), the centralspindlin complex and Kinesin-14/Ncd. We showed that 14-3-3 binds to the disordered region of Borealin, and this binding is regulated differentially by two phosphorylations on Borealin. Mutations at these two phospho-sites compromised normal Borealin localisation and centromere bi-orientation in oocytes, showing that phospho-regulation of 14-3-3 binding is important for Borealin localisation and function. The mass spectrometry data are available from ProteomeXchange, identifier <ID to be provided when available, PXD000xxx>.

**Author Summary:** Accurate segregation of chromosomes during cell division is fundamental for genome stability. Chromosome segregation is mediated by the spindle, which is made of dynamic microtubules and associated proteins that regulate microtubule behaviour. How these microtubule-associated proteins are regulated is not well understood. Furthermore, as oocytes have an exceptionally large volume of cytoplasm and lack centrosomes, regulation of microtubule-associated proteins is especially crucial for organisation and function of the meiotic spindle. In this study, we showed that 14-3-3, a protein that binds to phosphorylated proteins, plays an important role to regulate multiple microtubule-associated proteins in fly oocytes. The regulated proteins include subunits of the conserved kinase complex called the chromosomal passenger complex. We further found that interaction of one of the subunits with 14-3-3 is regulated by two phosphorylations, and that these phosphorylations are important for localisation and function of this subunit. As these proteins are widely conserved including humans, this study may provide an insight into chromosome mis-segregation in human oocytes, which is very frequent and a major cause of human infertility, miscarriages and congenital birth conditions.

## Introduction

Oocytes are specialised cells that undergo meiotic divisions to produce female gametes. Oocytes form the meiotic spindle to segregate chromosomes using mechanisms shared with mitotic cells, but also need to cope with unique challenges to assemble a spindle. The centrosomes are absent during meiotic spindle assembly in the oocytes of most animals, such as humans and *Drosophila melanogaster* [1], even though the centrosomes are the canonical microtubule organising centres in mitosis. Furthermore, oocytes have an exceptionally large volume of cytoplasm, but need to limit the spindle assembly to only the region around the chromosomes and not in other parts of the oocytes. A failure to suppress spindle assembly away from the chromosomes would lead to inefficient use of cellular resources, and most likely lead to aberrant cellular processes. Therefore, crucial proteins for spindle assembly are likely to be activated only around chromosomes but inactivated away from chromosomes.

The spatial regulation of spindle assembly in oocytes is not yet well understood, but one mechanism involving Ran and importin has been well described. As first shown in *Xenopus* egg extract, chromosome-associated RanGEF, Rcc1, converts Ran-GDP to Ran-GTP. Then Ran-GTP releases spindle assembly factors, such as TPX2, from inhibitory effects of importins to allow microtubule assembly around chromosomes (reviewed in [2]).

In living oocytes, there is mixed evidence for a role for Ran in chromosome-dependent spindle formation. On one hand, there is evidence that expression of dominant negative Ran impairs or delays meiosis I spindle assembly in human and mouse oocytes [3,4]. On the other hand, even when the Ran gradient was disrupted in a mouse oocyte by expressing hyperactive or dominant negative Ran, the oocyte still assembled a single spindle around chromosomes in meiosis I [5]. Similarly, in *Drosophila* oocytes, expression of dominant-negative or hyperactive Ran induced mild spindle defects in meiosis I but did not produce ectopic spindles [6]. These suggest that oocytes may have other pathways to spatially regulate spindle assembly, in addition to the Ran-importin pathway. The chromosomal passenger complex (CPC), containing Aurora B kinase, was proposed to provide an alternative chromosomal signal independently of Ran [7,8].

Recently, we proposed a new spatial regulation mechanism in *Drosophila* oocytes, involving the phospho-docking protein 14-3-3. 14-3-3 proteins are a well-conserved family of small, ubiquitous phospho-docking proteins involved in various cellular processes [9]. The 14-3-3 family is conserved across eukaryotes, with at least 7 different isoforms present in the human genome, while yeast, *Caenorhabditis elegans* and *Drosophila melanogaster* have 2 each (reviewed in [10,11]). 14-3-3 primarily exists as a dimer, either a homodimer of the same isoform or a heterodimer of two different isoforms.

Most of the described roles of 14-3-3 are through its direct binding to a target protein, with over 200 binding sites reported in the literature [12]. The majority of reported sites are centred on a phosphorylated serine or threonine, while in some rare cases 14-3-3 can bind in the absence of phosphorylation [13]. There are multiple mechanisms through which 14-3-3 can regulate its targets. For example, 14-3-3 can mask localisation motifs to regulate sequestration or shuttling of targets in or between subcellular compartments [14,15]. Some cases have also been reported of the 14-3-3 dimer acting as an adaptor, bringing together two proteins to increase their activity or interaction [16]. Additionally, Yaffe et al [17] propose that 14-3-3 might act as a “molecular anvil” to instigate conformational change in its binding partners. Finally, 14-3-3 might act as a chaperone of unfolded proteins, or facilitate clustering of proteins or complexes.

The importance of 14-3-3 in spindle formation in oocytes has already been demonstrated. 14-3-3 knockdown in *Drosophila* oocytes leads to defects in spindle bipolarity [18]. Similarly, depletion of 14-3-3η disrupts spindle assembly and polar body extrusion in mouse oocytes [19]. We previously identified a microtubule-crosslinking kinesin Ncd as a 14-3-3 target critical for spindle bipolarity in *Drosophila* oocytes [18]. Ncd and the mammalian orthologue HSET belong to the kinesin-14 family of minus-end directed microtubule motors [20], and both are required to focus the poles of the spindle in oocytes [21,22]. Our previous study proposed the mechanism by which 14-3-3 and Aurora B kinase allow Kinesin-14/Ncd binding of microtubules only near the chromosomes [18]. In the ooplasm away from chromosomes, 14-3-3 binds to Kinesin-14/Ncd at phosphorylated Serine 96, preventing Kinesin-14/Ncd from binding microtubules. In the vicinity of the chromatin, an additional phosphorylation nearby at Serine 94, by the chromatin-bound kinase Aurora B, inhibits 14-3-3 binding of kinesin-14/Ncd. This removes the inhibition, and specifically allows Kinesin-14/Ncd binding to spindle microtubules, promoting proper bipolar spindle formation.

Given the importance of spatial regulation for the oocyte, we ask if 14-3-3 regulates other important spindle-associated proteins in oocytes, and what mechanisms might be involved. By developing a new biochemical method, we identify various microtubule-associated proteins regulated by 14-3-3 in ovary extract. Among these proteins, of particular interest is the chromosomal passenger complex (CPC) subunit Borealin, because the CPC is one of the master regulators of cell division and not known to be regulated by 14-3-3 in any system. We show that Borealin is bound by 14-3-3, and this binding is differentially regulated by two phosphorylations. Mutations in these phospho-sites disrupts Borealin localisation to the spindle equator/centromeres and function in centromere bi-orientation. Collectively, these findings suggest that 14-3-3 plays a central role in a general regulatory system controlling microtubule-associated proteins (MAPs) in oocytes, analogous to the Ran-importin system.

## Results

### Identification of proteins regulated by 14-3-3 from *Drosophila* oocytes

We recently proposed a new mechanism of spatial regulation in oocytes, in which Kinesin-14/Ncd is regulated by the combined action of the phospho-docking protein 14-3-3 in the ooplasm and Aurora B kinase on chromosomes [18].

We hypothesise that this mechanism involving 14-3-3 and Aurora B provides a general way to spatially activate many spindle proteins only when near the chromosomes in oocytes. Here we tested this hypothesis by developing a new biochemical method to systematically identify proteins whose ability to binding microtubules is regulated by 14-3-3 in oocytes.

First, we have developed a novel method of purifying microtubule-associated proteins (MAPs) from *Drosophila* ovaries (**Fig 1A)** by adapting a previous method for embryos [23]. We took advantage of *Drosophila* ovaries being mainly made up of mature oocytes (arrested in metaphase I) by volume with a minimal contribution from mitotic cells. A hundred pairs of ovaries were dissected from mature flies and homogenised on ice to depolymerise microtubules. After the resulting lysate was cleared by ultracentrifugation, endogenous tubulin was polymerised by incubating with paclitaxel and GTP at room temperature. Microtubules, together with their associated proteins, were pelleted through sucrose cushion by ultracentrifugation. The pellet predominantly consists of tubulins and minimal amounts of other proteins (**S1 Fig**)

**Figure 1.**
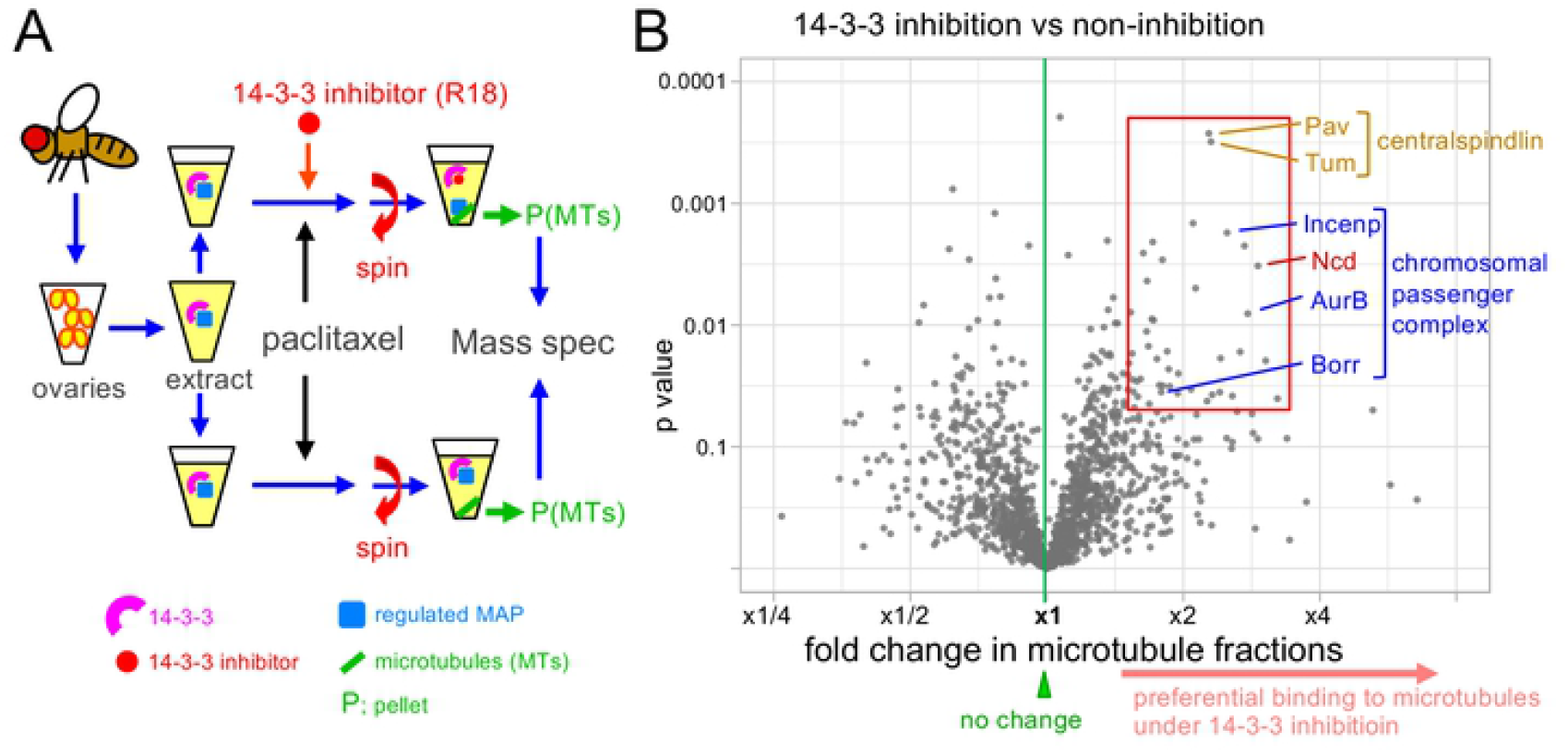
Identification of proteins whose microtubule binding is regulated by 14-3-3. (A) Dissected ovaries were homogenised and cleared by centrifugation to make soluble extract. It was separated into two and a 14-3-3 inhibitor (R18) was added to one. After microtubules are polymerised by addition of Taxol, they are sedimented by centrifugation through sucrose cushion. The pellets containing microtubules and their associated proteins were analysed by label-free mass-spectrometry. (B) Protein gel (C) Volcano plot showing the fold changes of the amounts of each protein detected in microtubule fraction in the presence of the 14-3-3 inhibitor in comparison to its absence on the X axis and the significance (p-value) on the Y axis. The red box contains 47 proteins that significantly increased their microtubule binding under 14-3-3 inhibition (the fold change ≥ 1.5 and the p ≤ 0.05).

Next we used a 14-3-3 inhibitor, R18. R18 is a synthetic peptide with 20 residues that has been shown to competitively bind to the 14-3-3 phospho-docking site with a higher affinity than native substrates [24]. This inhibitor has been shown to bind to various 14-3-3 isoforms across species [24–26], and therefore we expect it can inhibit both 14-3-3 isoforms in *Drosophila* oocytes. To identify MAPs regulated by 14-3-3 in oocytes, soluble ovary extract from wild type was divided into two aliquots. The 14-3-3 inhibitor R18 was added to one, and as a control, water was added to the other. After incubation with paclitaxel and GTP, microtubules and associated proteins were pelleted from both samples.

These two microtubule fractions were then analysed by LC-MS/MS for identification and label-free quantification.

### Quantitative mass-spectrometry has identified proteins potentially regulated by 14-3-3

To determine the relative amounts of each protein in the microtubule fraction with or without 14-3-3 inhibition, six pairs of biological replicates were analysed by label-free quantification using LC-MS/MS (**S2-7 Table**). As a result, we have detected 3,564 proteins in ovaries, including 24 out of 32 known spindle MAPs in our dataset. Among them, we were able to quantify 1,504 proteins (42%) in at least 2 out of the 6 experiments (**S1 Table**). These proteins were plotted as a volcano plot with fold differences on the X axis and the confidence level (p-value) on the Y axis (**Fig 1B; S1 Table**). In this plot, proteins on the top right increased their binding to microtubules under 14-3-3 inhibition. In total, 47 proteins were detected in microtubule fractions at a higher abundance in the presence of the 14-3-3 inhibitor than in its absence with a good confidence (defined by p<0.05 and the ratio>1.5; **Fig 1B**). Therefore 14-3-3 is likely to suppress microtubule binding of these proteins in oocytes.

It is possible that some identified proteins are not regulated directly by 14-3-3, and instead form a complex with a protein regulated by 14-3-3. To test this possibility, we examined whether any of these 14-3-3 regulated proteins or their orthologues are known to physically interact with each other using STRING database ([27]; **Fig 2A**). We found 48 known protein-protein interactions among these 47 proteins, which is significantly higher than expected from a random set of 47 proteins (17 interactions; p=1.2e^-09^). This is consistent with the possibility that microtubule binding of some proteins is indirectly regulated by 14-3-3 through interactions with other proteins.

**Figure 2.**
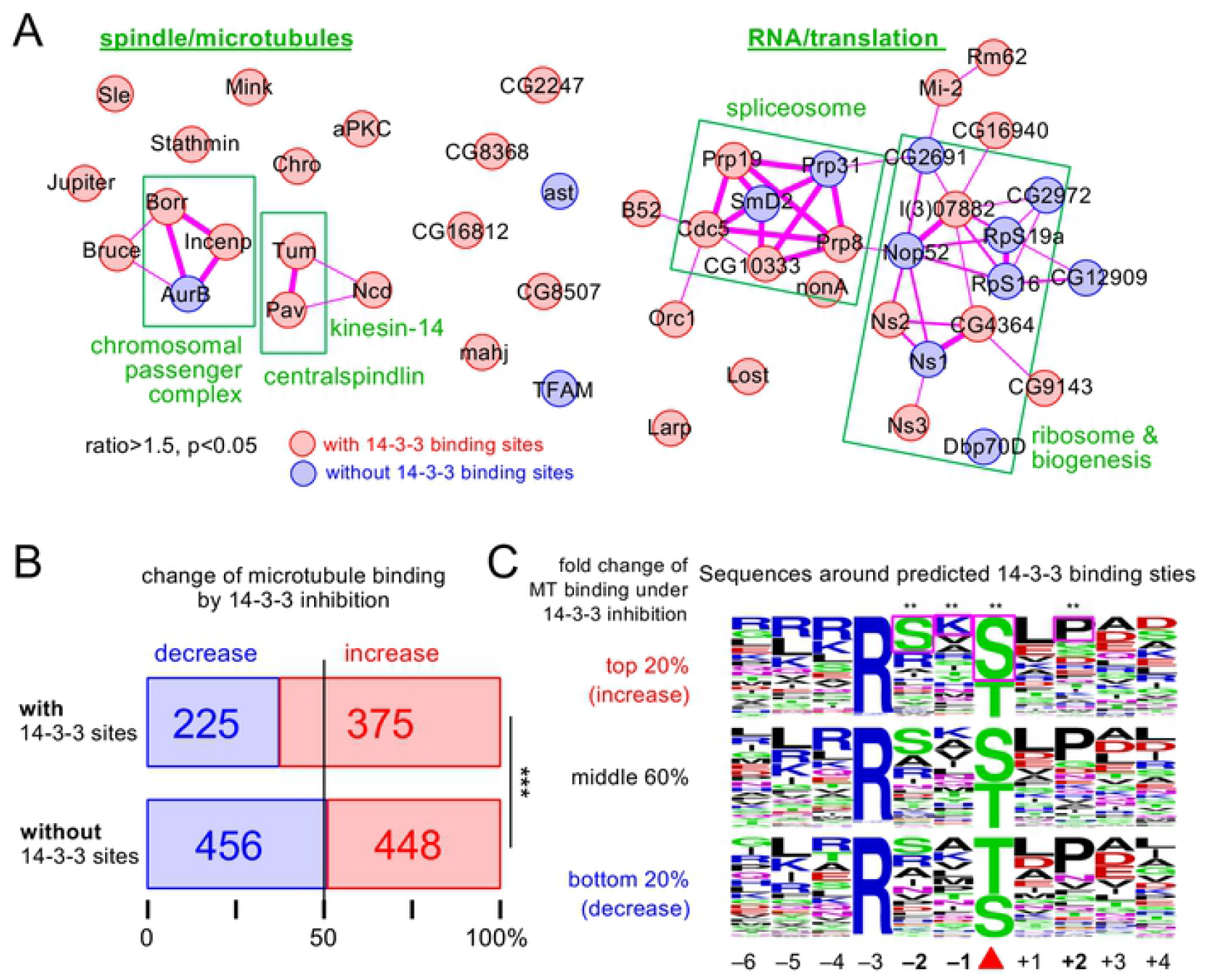
Bioinformatics analysis. (A) Physical protein-protein interactions known among 47 proteins that significantly increased their microtubule binding under 14-3-3 inhibition (the fold change ≥1.5 and the p ≤0.05; the boxed area in Fig 1C). Red and blue indicate proteins with at least one predicted 14-3-3 binding sites and without them, respectively. Thickness of lines indicate confidence levels of evidence. (B) The numbers and proportions of proteins with or without predicted 14-3-3 sites that increased or decreased in the microtubule fractions by 14-3-3 inhibition. (C) Sequences around predicted 14-3-3 binding sties among proteins with top 20%, middle 60% and bottom 20% of all detected proteins in the order of the fold change in microtubule fraction in the presence of the 14-3-3 inhibitor. Coloured boxes indicate the residues whose frequencies are significantly different among the top 20% than the frequencies among the rest of proteins. ** indicates p<0.01.

One of the proteins that increased binding to microtubules under 14-3-3 inhibition with a high degree and confidence was a known 14-3-3 regulated protein in oocytes, Kinesin-14/Ncd (**Fig 1B**). In addition, both subunits of centralspindlin (MKlp1/Pav, RacGAP/Tum) increased their binding to microtubules under 14-3-3 inhibition (**Fig 1B, 2A**). This is consistent with the previous study that shows MKlp1/Pav is directly regulated by 14-3-3 in human mitotic cells [28], although this has not been shown in oocytes. These results confirmed that our new method can successfully identify 14-3-3 regulated proteins. Similarly, three out of the four CPC subunits (Aurora B, Incenp and Borealin) significantly increased microtubule binding under 14-3-3 inhibition, except the smallest subunit Survivin/Deterin (**Fig 1B, 2A**). The CPC is not yet known to be regulated by 14-3-3 in any organism, and therefore was investigated further.

A broader question is whether 14-3-3 regulates only a few microtubule-associated proteins or many proteins in oocytes. To gain an insight into this question, we tested whether the presence of a predicted 14-3-3 binding sites correlated with either an increase or decrease in microtubule binding under 14-3-3 inhibition (**Fig 2A**). If any changes under 14-3-3 inhibition are purely due to random statistical variations rather than 14-3-3 regulation, the number of proteins which increases or decreases in microtubule binding would be 50:50, regardless of whether proteins can bind 14-3-3. Among 1,504 proteins quantifiable in our experiments, 600 proteins had at least one site predicted at a high confidence to bind 14-3-3. Interestingly, among these 600 proteins, 375 increased (62%) and 225 decreased (38%) microtubule binding under 14-3-3 inhibition, showing proteins with predicted 14-3-3 binding sites are much more likely to increase microtubule binding (a difference of 150 proteins; p<0.001; **Fig 2B**). In contrast, among the remaining 904 proteins without predicted 14-3-3 binding sites, 448 increased (49%) and 456 decreased (51%) microtubule binding, which is not significantly different from 50:50 (p=0.82; **Fig 2B**). This suggests that 14-3-3 potentially affects microtubule binding of many proteins in oocytes, and more often suppresses their microtubule binding than promotes it.

In the case of Kinesin-14/Ncd, 14-3-3 binding is regulated by phosphorylation at two sites, and regulating 14-3-3 binding is important for spatially controlling the microtubule binding activity of Kinesin-14/Ncd in oocytes. If 14-3-3 binding to a subset of proteins is regulated by a common mechanism such as phosphorylation by the same kinases, we may see a bias in sequences surrounding 14-3-3 binding sites in addition to the 14-3-3 binding motif. To gain an insight into regulation of 14-3-3 binding, sequences surrounding predicted 14-3-3 binding sites were compared between proteins whose microtubule binding is, or is not, regulated by 14-3-3. All identified proteins were divided into 5 bins according to the fold change in microtubule association under 14-3-3 inhibition. Sequences around predicted 14-3-3 sites on proteins in each bin were pooled together. As the three middle bins are similar, the top 20% (increase under inhibition), the middle 60% and the bottom 20% (decrease under inhibition) were compared (**Fig 2C; S8 Table**). Serine at −2, serine at 0 and lysine at −1 are significantly overrepresented among the top 20%, while proline at +2 is underrepresented. As previously shown for Kinesin-14/Ncd, Serine at −2 can be phosphorylated by a kinase and this would negatively regulate 14-3-3 binding. Overrepresentation of serine over threonine at 0 (14-3-3 binding site) might also be interesting, as the phosphatase PP2A-B55 is known to prefer phospho-threonine over phospho-serine for dephosphorylation [29].

### The disordered region of Borealin physically interacts with 14-3-3 in a phospho-dependent manner

Our results showed that three out of the four CPC subunits (Aurora B, Incenp and Borealin) significantly increased their quantities in microtubule fractions under 14-3-3 inhibition (**Fig 1C**). Previous studies from us and others showed that the CPC has crucial roles in oocytes, including spindle microtubule assembly, spindle organisation, and bi-orientation of homologous centromeres [30–35]. The CPC dynamically localises to chromosomes, centromeres and the spindle equator in oocytes [33,36]. However, the CPC has not been reported to be regulated by 14-3-3 in any organism.

Among the CPC subunits, we identified 3 potential sites in Borealin and Incenp that match a consensus 14-3-3 binding sequence (**Fig 3A)** by bioinformatic analysis (Madeira et al., 2015). Among them, one site on Borealin (S163) not only matches the 14-3-3 binding consensus, but also shares a high similarity with the 14-3-3 binding sites of Kinesin-14/Ncd and Kinesin-6/MKlp1/Pav (**Fig 3B**). A distinct feature of these sites is the serine residue at the −2 position (S161) that matches the Aurora B phosphorylation consensus (R/KxS/T).

**Figure 3.**
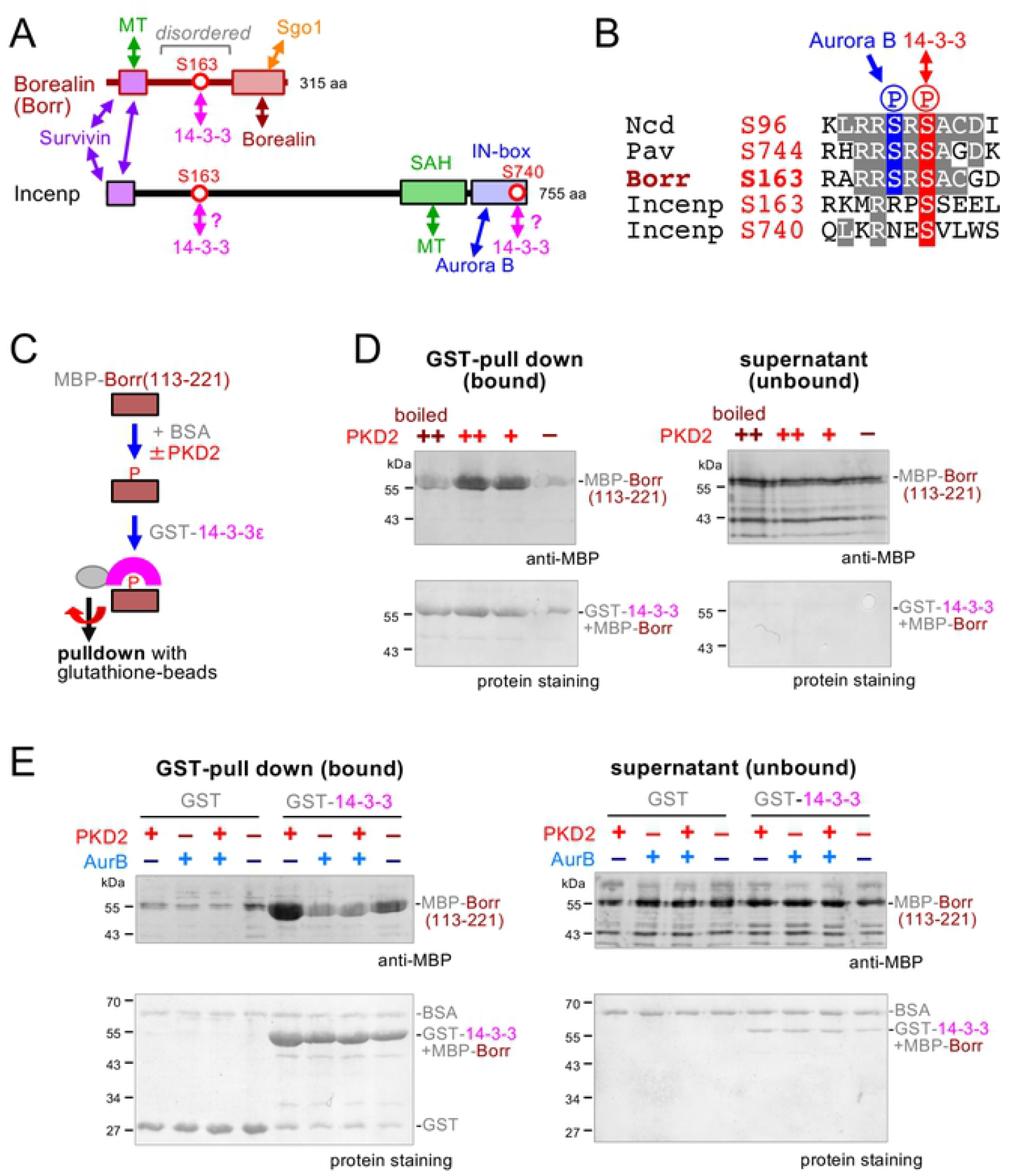
14-3-3 binds to the disordered region of Borealin and regulated by two phosphorylations. (A) Diagram of domain organisation of Borealin and Incenp proteins. Known protein-protein interactions are indicated as double arrows. Three predicted 14-3-3 binding sites (red circles; Borealin S163 and Incenp S163 and S740) are identified among the CPC subunits. Borealin S163 is located within an intrinsically disordered region between domains interacting with the CPC subunits. MT; microtubule. (B) Sequences surrounding predicted 14-3-3 binding sites. The sequence surounding Borealin has high similarity to Ncd S96 and Pav S744, containing serine at −2 position that is potentially phosphorylated by Aurora B kinase. (C) Testing interaction between the phospho-docking protein 14-3-3 and the disordered region of Borealin (Borr). Purified MBP-tagged Borealin fragment was incubated with or without PKD2, together with BSA. GST-14-3-3ε was added and pulled down by centrifugation using glutathione beads. Proteins bound to the beads and unbound fraction (supernatant) were analysed by western blotting using an MBP antibody. (D) MBP-Borealin(113-221) interacts with GST-14-3-3 in a phospho-dependent manner. MBP-Borealin(113-221) was incubated with two different concentrations of human PKD2 kinase (++, higher; +, lower), with the higher concentration of inactivated PKD2 (“boiled ++”) or without PDK2 (–). A western blot of proteins bound to the glutathione beads and unbound fractions (supernatants) was carried out using an MBP antibody and total protein staining. (E) An additional phosphorylation by Aurora B prevents PKD2-phosphorylated MBP-Borealin(113-221) from interacting with GST-14-3-3. Borealin(113-221) was incubated with human PKD2 kinase alone, human Aurora B kinase alone, both kinases or without kinases, and tested for pull down using GST or GST-14-3-3ε. Western blots of bound or unbound proteins were analysed by a western blot using an MBP antibody or total protein staining. PKD2-phosphorylated MBP-Borealin(113-221) specifically interacted with GST-14-3-3ε, while MBP-Borealin(113-221) doubly phosphorylated by PKD2 and Aurora B did not.

This potential 14-3-3 binding site (S163) on Borealin is located in a disordered region between the Incenp/Survivin interaction domain and the Borealin dimerisation domain [37,38] (**Fig 3A**). To experimentally test whether 14-3-3 can bind this disordered region of Borealin, the region (residues 113-221) was produced with MBP-tag in bacteria. After purification (**S2 Fig**), MBP-Borealin(113-221) was phosphorylated by incubation with human PKD2 kinase at two different concentrations. PKD2 was used as it is known to efficiently phosphorylate the similar 14-3-3 binding site in Ncd *in vitro* [18]. As negative controls, MBP-Borealin(113-221) was either incubated with PKD2 inactivated by boiling or incubated without the kinase. After the reaction, MBP-Borealin(113-221) was incubated with bacterially produced GST-14-3-3 (**S2 Fig**). GST-14-3-3 was then pulled down by glutathione-beads and analysed by western blot using an MBP antibody (**Fig 3C**). We found that MBP-Borealin(113-221) was efficiently pulled down with GST-14-3-3 only after treatment with active PKD2 (**Fig 3D**). This indicates that this disordered region of Borealin binds to 14-3-3 in a phospho-dependent manner.

Next, we tested whether an additional phosphorylation by Aurora B can prevent Borealin from binding to 14-3-3, as a potential Aurora B phosphorylation site (S161) is located near the predicted 14-3-3 binding site (S163). MBP-Borealin(113-221) was first incubated with PKD2, Aurora B, both kinases or no kinases, and then tested for 14-3-3 binding by pulling down with GST-14-3-3. We found that MBP-Borealin(113-221) phosphorylated by both PKD2 and Aurora B bound poorly to 14-3-3, much less than when phosphorylated by PKD2 alone (**Fig 3E**). Therefore, the additional phosphorylation by Aurora B prevents binding of 14-3-3 to PKD2-phosphorylated MBP-Borealin(113-221).

### Phospho-regulation of 14-3-3 binding is important for efficient Borealin localisation

Our biochemical analysis showed that 14-3-3 binding to Borealin is differentially regulated by two phosphorylations (**Fig 3B**,**E**). To define the *in vivo* role for this phospho-regulation in oocytes, we generated GFP-tagged versions of two non-phosphorylatable mutants at 14-3-3 binding site (S163A) and the Aurora B phosphorylation site (S161A).

Their localisation was examined in the absence of the endogenous protein by expressing the GFP-tagged RNAi-resistant wild-type and non-phosphorylatable mutants together with a short hairpin RNA (shRNA) against the endogenous gene, and the chromosomal marker Rcc1-mCherry in oocytes. Live imaging showed that wild-type Borealin localised to the spindle equator and centromeres (**Fig 4A; S9 Table**), as reported previously for the CPC subunits [33,39].

**Figure 4.**
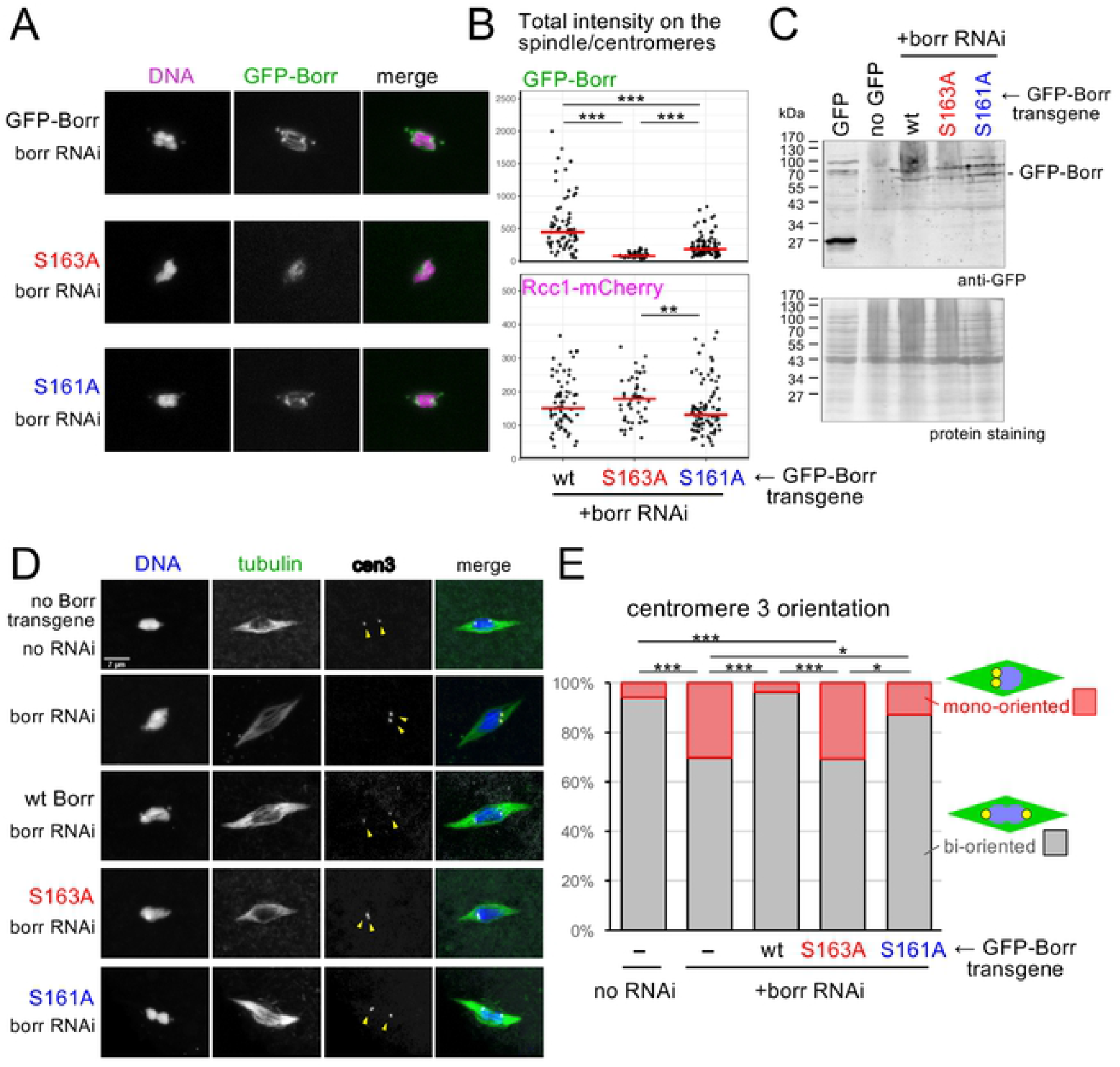
Non-phosphorylatable mutations compromises the localisation of Borealin and bi-orientation of centromeres in oocytes. (A) Non-phosphorylatable mutations (S163A, S161A) reduce the Borealin localisation to the spindle and centromeres. Fluorescence was observed in live oocytes expressing the GFP-tagged RNAi-resistant wild-type or non-phosphorylatable mutants together with a short hairpin RNA (shRNA) against the endogenous gene and a red chromosomal marker, Rcc1-mCherry. The images were presented using the same condition of capture and contrast adjustment for comparison. Arrowheads indicate the position of centromeres. (B) The total signals of GFP-Borealin and Rcc1-mCherry on the spindle and centromeres above the background was quantified. Red lines indicate the median signal intensities. *** and ** indicate p<0.001 and p<0.01, respectively (Wilcoxon rank sum test). (C) Western blot of ovaries expressing GFP, wild-type ovaries without expression of GFP, and ovaries expressing wild-type or mutant GFP-Borr (S163A or S161A) in the background of *borr* RNAi, probed with an anti-GFP antibody and protein staining. (D) Non-phosphorylatable mutations (S163A, S161A) compromises the Borealin function in the bi-orientation of centromeres. α-tubulin and peri-centromere 3 (dodecasatellite; arrowheads) were visualised by Immunostaining combined with *in situ* hybridisation in *borr* RNAi oocytes with or without expression of the wild-type or non-phosphorylatable Borealin mutants. (E) Frequencies of bi-orientation of homologous centromeres of chromosome 3. Two separate dodecasatelite signals near both ends of the chromosome mass are considered as bi-oriented centromeres, and one or closely located two signals on the one side of the chromosome mass are considered as mono-oriented centromeres. *** and * indicate p<0.001 and p<0.05, respectively (Fisher exact test).

A non-phosphorylatable Borealin mutant at the 14-3-3 binding site (S163A) nearly abolished the localisation on the spindle equator and centromeres (**Fig 4A**). The non-phosphorylatable Borealin mutant at the Aurora B phosphorylation site (S161A) still localised, but with a reduced intensity, on the spindle equator and centromeres in comparison to wild-type Borealin (**Fig 4A**). For quantification, a total intensity of GFP signals including both the spindle equator and centromeres above the background were measured. The S163A mutation resulted in a dramatic drop in the GFP signal intensity by 82% in comparison to wild-type Borealin (p<0.001), while the S161A mutation resulted in a less dramatic but significant reduction (by 59%; p<0.001) (**Fig 4B**). In contrast, intensities of the chromosome signal (Rcc1-mCherry) were comparable, suggesting that a difference of the Borealin signal intensity is due to the reduction in the localisation (**Fig 4B**), rather than a secondary effect. GFP-Borealin mutant proteins were expressed in oocytes at comparable levels to wild-type GFP-Borealin (**Fig 4C**).

These results revealed that phospho-regulation of 14-3-3 binding to Borealin is important for Borealin localisation in oocytes. The phospho-dependent 14-3-3 binding is crucial for Borealin localisation, while the phosphorylation by Aurora B is less crucial but still important.

### 14-3-3 binding to Borealin is important for bi-orienting homologous centromeres

To establish the functional significance of the phospho-regulation of 14-3-3 binding in oocytes, we further examined the oocytes expressing the wild-type or mutant Borealin transgene in a background in which the endogenous gene is silenced. We then examined spindle morphology and centromere positions in mostly metaphase I arrested oocytes by immunostaining tubulin combined with *in situ* hybridisation using a peri-centromere probe specific to chromosome 3 (dodecasatellite; [40]).

We examined oocytes expressing an shRNA against Borealin and without Borealin transgenes alongside a control without shRNA expression (**Fig 4D,E**). In the control without shRNA expression, one centromere 3 signal was found on each side of the chromosome mass in nearly all (95%) of oocytes, showing a pair of homologous centromeres which are bi-oriented and pulled apart to both poles. Only 5% of oocytes showed a mono-orientation in the control. In contrast, 30% of RNAi oocytes showed mono-orientation (p<0.001). The spindle morphologies look similar between Borealin RNAi and the control. This may either reflect partial depletion of Borealin or indicate that Borealin is not essential for normal spindle morphology.

Expression of GFP-tagged wild-type Borealin fully rescued the centromere bi-orientation defect of Borealin RNAi, and the frequency of mono-orientation went back to a similar level to oocytes without RNAi. In contrast, expression of GFP-tagged Borealin(S163A) failed to rescue the defect, and the frequency of mono-orientation stayed at a similar level to the RNAi oocytes without a transgene. This is consistent with the large reduction in localisation to the spindle/centromeres we observed above (**Fig 4D,E; S10 Table**). Expression of GFP-tagged Borealin(S161A) showed an intermediate effect, which is also consistent with partial loss of localisation. These results showed that phosphorylation for 14-3-3 binding at S163 is essential for Borealin function, while phosphorylation of the flanking site (S161) is not essential but appears to play a role. These results showed that phospho-regulation of 14-3-3 binding is important for the centromere bi-orientation function of Borealin.

## Discussion

By microtubule co-sedimentation combined with quantitative mass-spectrometry, we identified various proteins whose microtubule binding is regulated by 14-3-3 in *Drosophila* oocytes. Among them, we showed that 14-3-3 binds to Borealin, a subunit of the chromosomal passenger complex (CPC). Two phosphorylations regulating this 14-3-3 binding to Borealin are important for Borealin localisation and centromere bi-orientation in oocytes.

Using a newly developed method, we quantified the relative amounts of over 3,000 proteins associated with microtubules in ovary extract with or without 14-3-3 inhibition. Our analysis showed that 14-3-3 suppresses microtubule binding of a substantial number of proteins in ovaries. Among these proteins, Kinesin-14/Ncd and the centralspindlin subunits were previously reported to be regulated by 14-3-3 [18,28] in *Drosophila* oocytes and human cells, respectively. We also identified three subunits of the CPC, one of the master regulators of cell division [32,41]. To our knowledge, this is the first report to show that the CPC is regulated by 14-3-3.

We found that the CPC subunit Borealin has a predicted 14-3-3 binding site (S163) in its disordered middle region. The surrounding sequence shares a striking similarity to the 14-3-3 binding sites identified in Kinesin-14/Ncd and the centralspindlin subunit MKlp1/Pavarotti [18,28]. Through *in vitro* studies, we showed that the disordered middle region of Borealin binds to 14-3-3 in a phospho-dependent manner. Additional phosphorylation by Aurora B reverses the binding, in a similar manner to that reported for Kinesin-14/Ncd [18]. A non-phosphorylatable mutation predicted to abolish 14-3-3 binding (S163A) dramatically reduced the amount of Borealin at the spindle and centromeres. This mutation also abolishes the function of Borealin in bi-orienting homologous centromeres. On the other hand, a non-phosphorylatable mutation of the predicted Aurora B site (S161A) also compromises the normal localisation and function of Borealin but not abolishing them. This partial requirement of the Aurora B phosphorylation can be explained if 14-3-3 binding to Borealin is prevented by redundant action of two enzymes: a phosphatase dephosphorylating the 14-3-3 binding site (S163) and Aurora B phosphorylating the neighbouring site (S161). Similar effects were also observed in the equivalent mutations in Kinesin-14/Ncd [18], strengthening our hypothesis that a common mechanism regulates Borealin and Kinesin-14/Ncd.

Based on the model proposed by Beaven et al [18] for Ncd, we propose that Borealin is prevented from binding microtubules in the oocyte cytoplasm by its association with 14-3-3. When it is in the vicinity of the chromatin, Aurora B kinase activity reverses the 14-3-3 binding and allows the CPC to bind spindle microtubules. We showed that this regulation is important for CPC’s role in bi-orienting homologous centromeres. In our model, spatial regulation mediated by 14-3-3 and Aurora B allows Borealin to bind selectively to spindle microtubules. Abolishing this spatial regulation would dramatically reduce the effective concentration of Borealin near the spindles by binding to numerous non-spindle microtubules in the large volume of the oocytes.

This study has established 14-3-3 as a general regulator of microtubule-associated proteins (MAPs). 14-3-3 may play an important role in spatially regulating many MAPs in oocytes in combination with other regulators such as Aurora B or phosphatases, in parallel to the Ran/importin system. Future studies of these 14-3-3 regulated proteins will shed light on how microtubules are regulated in oocytes. The study has also revealed a novel regulation of the CPC. The CPC is one of the conserved master regulators of cell division, and dynamically localises to chromosomes, centromeres and spindle microtubules. Although the chromosome/centromere localisation of the CPC has been intensely studied, microtubule binding and its regulation are paid much less attention. Our study identified a novel mechanism regulating microtubule binding activity of the CPC, which is essential for bi-orienting homologous chromosomes in oocytes. As the CPC is known to have other microtubule binding domains than this disordered region of Borealin [42], further studies are required to establish whether this region of Borealin defines a new microtubule binding domain or 14-3-3 binding to this region interferes with microtubule binding of other domains.

## Materials and Methods

### Fly maturation and generation of GFP-Borr flies

Standard fly techniques were followed [43]. *Drosophila melanogaster* stocks were cultured on standard cornmeal medium at 25°C. *w*^*1118*^ was used as wild-type. Less than 1 day old female adults were matured for 3 days at 25°C in the presence of males before dissection of ovaries.

To generate Borealin transgenes, the full-length Borr open reading frame was amplified from LD36125 by PCR using PrimeStar (Takara) and cloned between AscI and NotI sites of pENTR (ThermoFisher) using Gibson assembly (HiFi; NEB). To make the gene resistant to RNAi, silent mutations (from GCG GTG TTC to GCC GTC TTT) were introduced by PCR using primers containing the mutations and followed by Gibson assembly. Further mutations (TCC to GCC for S163A; AGT to GCC for S161A) were introduced using the same method. The absence of unwanted mutations were confirmed by Sanger sequencing. These entry clones were recombined with the destination vector φMGW [36] by LR Gateway Clonase II (ThermoFisher).

Transgenic flies expressing RNAi-resistant GFP-Borr (wild-type/S163A/S161A) under the maternal α-tubulin promoter were generated using φC31 integrase-mediated transgenesis at the *VK18* site [44] on the second chromosome, performed by BestGene Inc. Each transgene of GFP-Borr or variant was then recombined with another transgene at the *attP40* site (Markstein et al, 2008) expressing short hairpin (sh) RNA against *borr* (HMC04381; [45]; 55942 from Bloomington Drosophila Stock Center).

### Microtubule cosedimentation with the 14-3-3 inhibitor R18

Microtubules and their associated MAPs were purified from ovaries by modifying protocols previously used in embryos [46] and in S2 cells [23]. Ovaries from matured female *w*^*1118*^ flies were dissected in 1x BRB80 buffer (80 mM PIPES-KOH pH 6.8, 1 mM MgCl_2_, 1 mM Na_3_EGTA) supplemented with phosphatase inhibitors (1mM DTT (Promega), 1mM PMSF, Complete EDTA-free Protease Inhibitor Mixture Tablets MINI, diluted according to manufacturer instructions (Roche), 15 mM Na_3_VO_4_, 10 mM p-nitrophenyl phosphate, 1 μM okadaic acid). Ovaries were snap frozen in liquid nitrogen and stored at −80°C. About 100 pairs of ovaries (∼150 mg in wet weight) were defrosted and pooled. An equal volume of 2x BRB80+phosphatase inhibitors was added. Ovaries were homogenised in a pre-cooled Dounce homogeniser and incubated on ice for 30 minutes to depolymerise microtubules, then cleared by centrifugation at 13,000 rpm/13,628 x g for 15 minutes at 4°C. The supernatant was cleared again by 2 rounds of ultracentrifugation at 100,000 rpm (358,400 x g; Beckman TLA-120.2) for 10 minutes at 0°C, and then incubated at 30°C for 20 minutes, followed by ultracentrifugation at 100,000 rpm for 10 minutes at 22°C. Actin was depolymerised by the addition of 1 µg/mL Latrunculin A (Cambridge Bioscience), 2 µg/mL Latrunculin B (Cambridge Bioscience), 0.1 mM DTT (Promega) and 1 mM GTP (Sigma), and an aliquot of the lysate was kept for analysis (‘input’).

The remaining supernatant was split into two portions, one of which (‘sample’) was treated with R18 peptide (R18 trifluoroacetate, Sigma) at a final concentration of 100 µM to inhibit 14-3-3 activity. To the other (‘control’) was added an equal volume of water. Both were then incubated for 5 minutes at room temperature. Paclitaxel (Sigma-Aldrich) was subsequently added to both sample and control at a final concentration of 20 μM, and incubated for 30 minutes to allow microtubules to polymerise. Microtubules were pelleted twice through 50% sucrose cushions containing 20 μM paclitaxel via ultracentrifugation at 50,000 × g for 35 minutes at 22°C. Pellets were resuspended in 50 μl of 1x BRB80 + phosphatase inhibitors supplemented with 20 μM paclitaxel. An equal volume of 3x SDS sample buffer + 5% β-mercaptoethanol was added to samples before boiling at 95°C for 2 minutes to denature proteins. Samples were stored at −20°C. Aliquots of samples were analysed by western blot. Total proteins on the membrane were stained with MemCode reversible staining kit (Thermo-Fisher) first, and then with a rat polyclonal antibody against α-tubulin (1:500) followed by IRDye 800CW conjugated goat anti-mouse IgG antibody (LI-COR). The signals were detected with an Odyssey CLx imaging scanner (LI-COR). The brightness and contrast were adjusted uniformly across the entire area in a linear manner without removing or altering features.

### Mass spectrometry

Pellets from microtubule cosedimentation were separated using a NuPAGE 12% Bis-Tris gel with MOPS running buffer and stained with Colloidal Blue Staining Kit (Invitrogen). 2 lanes were run for each pellet. The gel region containing the tubulin band was excised and discarded from both sample and control lanes. The remaining gel regions were trypsin-digested and reduced/alkylated using standard procedures [47]. Following digestion, samples were diluted with an equal volume of 0.1% TFA and spun onto StageTips as described by Rappsilber et al. [48]. Peptides were eluted in 80% acetonitrile in 0.1% TFA and concentrated 40x by vacuum centrifugation.

Samples were prepared for LC-MS/MS analysis by diluting them to 5 μL with 0.1% TFA. MS-analysis was performed on an Orbitrap™ Fusion™ Lumos™ tribrid™ mass spectrometer (Thermo Fisher Scientific, UK) coupled on-line to Ultimate 3000 RSLCnano Systems (Dionex, Thermo Fisher Scientific, UK). Peptides were separated by a PepMapTM RSLC C18 EasySpray column (2 µm, 100Å, 75 µm x 50 cm) (Thermo Fisher Scientific, UK), operating at 50°C. Parameters are described in **S11 Table**. The peptide gradient was: 2 to 40% buffer B in 140 min, then to 95% in 11 min. The percentage of buffer B remained at 95 for 5 minutes and returned back at 2 one minute after. Peptides were selected and fragmented by higher-collisional energy dissociation (HCD) [49] with normalised collision energy of 30. Raw files were processed using MaxQuant software platform [50] version 1.5.2.8 against the complete *Drosophila melanogaster* proteome (Uniprot, released in September 2016), using Andromeda [51]. Raw mass spectrometry proteomics data were deposited to the ProteomeXchange Consortium (http://proteomecentral.proteomexchange.org) via the PRIDE partner repository [52] with the dataset identifier <ID to be provided when available, PXD000xxx>.

### Bioinformatic analysis

Peptides were assigned to Uniprot accession numbers, and arranged as one accession per row. When more than one accession was assigned per gene, the accession with the lower score was removed. Accessions assigned only one peptide cannot be confidently identified and so were removed. Accessions denoting different RNA transcripts from the same gene were collapsed into one accession, again keeping the highest scored accession in the event of duplicates. The ratio between sample and control intensities for each gene was calculated as log_2_(sample) – log_2_(control). When either the sample or control intensity was 0, these accessions were removed from the graph and the statistical analysis and analysed separately.

To determine the consistency between replicates, log_2_(sample) and log_2_(control) value distributions were compared between replicates, as well as the line of best fit for the graph of log_2_(sample) vs log_2_(control). 6 out of 8 replicates were determined to be similar enough to continue with analysis. To compare the sample and control intensities for each accession, the given accession had to be detected in at least two replicates for both sample and control. The ratios were calculated as above, and the p-value for each ratio calculated using a two-tailed unpaired t-test. Then the ratio and p-value for each protein were plotted in log_2_ as a volcano plot (**Fig 2C**).

Gene ontology terms were downloaded from Flybase [53]. Data manipulation and statistical analyses were performed using Excel (Microsoft) and RStudio (RStudio Team, 2020) using the R scripting language [54]. Graphs were plotted using ggplot2 [55]. Candidate 14-3-3 binding sites were predicted using the 14-3-3 Pred tool [56] on the basis of a consensus score above 0.9 and individual scores above the default thresholds (ANN 0.55; PSSM 0.80; SVM 0.25) for all 3 prediction methods. Likely kinase target sequences were predicted in Borealin sequence using the GPS 3.0 programme [57] with medium threshold. Protein fasta sequences were obtained from UniProt. Protein interaction data for network analysis was downloaded from STRING-db [27] on 29 Nov 2021. Included interactions were limited to physical subnetwork above medium confidence (0.400) sourced from experiments and databases. 14-3-3 site sequence comparison analysed and presented using Weblogo (https://weblogo.berkeley.edu/, [58]).

### GST-14-3-3 binding to MBP-Borr including phosphorylation

To generate a construct expressing MBP-Borr(113-221) in *E. coli*, Borr open reading frame corresponding to amino acids 113-221 was first amplified with a stop codon from LD36125 by PCR using PrimeStar (Takara) and cloned between AscI and NotI sites of pENTR (ThermoFisher) using Gibson assembly (HiFi; NEB). This entry clone was recombined using LR clonase II (ThermoFisher) with a destination vector modified from pMAL-c2 (NEB) by insertion of the Gateway cassette (ThermoFisher) at the XmnI site. GST-14-3-3ε was produced using a construct previously reported [18].

To test if Borealin is bound by 14-3-3, MBP-Borr(113-221) and GST-14-3-3ε were purified from *E. coli* (BL21/pLysS) carrying the expression constructs using amylose beads (NEB) and S-glutathione beads (GE Healthcare), respectively. Fractions were eluted using 10 mM reduced glutathione through a polypropylene column (Qiagen), and those with the highest concentrations combined. Combined elutes were then dialysed overnight using Slide-a-Lyser dialysis cassettes (Thermo) in 1 l of dialysis buffer (50 mM Na-phosphate buffer pH7.6, 250 mM KCl, 1 mM MgCl_2_, 5 mM β-mercaptoethanol). Subsequently, the dialysed purified GST-14-3-3ε was concentrated to 500 μl total using Ultra-4 Spin Columns (Amicon), before being aliquoted, supplemented with 10% final concentration of glycerol, snap frozen in liquid nitrogen and stored at −70°C.

After thawing on ice, 3 μg (3.75 μl) of purified MBP-Borr(113-221) protein were cleared of aggregates by centrifugation for 1 minute at 14,000 rpm (13,628 x g), and incubated with 20 μl washed S-glutathione beads for 30 minutes at 4°C. After the beads were removed by centrifugation, 7.5 μg of MBP-Borr(113-221) was incubated with either 450 ng Aurora B (Cambridge Biosciences), 337.5 ng of PKD2 (Cambridge Biosciences), both Aurora B and PKD2 or no kinase. Phosphorylation was carried out in a two-step incubation at 30°C in phosphorylation buffer (20 mM HEPES pH 7.4, 2 mM MgCl_2_, 1 mM ATP, 27.5 mM KCl, 1 mM DTT, 0.2 mg/mL BSA, 1 mM EGTA), with Aurora B added for the first 30 minutes, followed by addition of PKD2 and a further 60 minutes at 30°C. Phosphorylated fragments (5 μl) were then mixed with 20 μg GST or GST-14-3-3 in 500 μl Pulldown buffer (25 mM Tris-Cl pH=7.6, 150 mM NaCl, 0.5% Triton X-100, 0.3 mM Na_3_VO_4_) and allowed to bind for 30 minutes on ice. They were then incubated with 20 μl washed S-glutathione beads on a rotator at 4°C for 1 hour. The supernatant/unbound fraction was removed and kept for analysis, and beads were then washed 3 times in 800 μl of Pulldown buffer. 2x SDS loading buffer was added to beads and supernatant samples at 1:1 ratio, supplemented with 5% final volume of β-mercaptoethanol, and were boiled for 2 minutes at 95°C, before analysis by western blot. Proteins were separated by SDS-PAGE and transferred onto a nitrocellulose membrane using the Mini-PROTEAN and Mini Trans-Blot system (Biorad). Total proteins on the membrane were stained with MemCode reversible staining kit (Thermo-Fisher). After being destained, the membrane was incubated with a rat polyclonal antibody against MBP (1:500) followed by IRDye 800CW conjugated goat anti-rat IgG antibody (LI-COR). The signals were detected with an Odyssey CLx imaging scanner (LI-COR). The brightness and contrast were adjusted uniformly across the entire area in a linear manner without removing or altering features.

### Live-imaging of oocytes

To visualise the localisation of GFP-Borr in live oocytes, flies carrying two transgenes that express GFP-tagged Borr (wild-type, S163A or S161A) under the maternal α-tubulin promoter and shRNA against *borr* under the UASp promoter were crossed with flies carrying *GAL4-VP16* (*V37*; Bloomington Drosophila Stock Centre 7063) and *Rcc1-mCherry* [36] both under the maternal α-tubulin promoter. Ovaries from 3-day matured flies were dissected one at a time in Halocarbon oil (700; Halocarbon) on a cover slip. Stage 14 oocytes were identified by their morphology, namely a well-formed chorion and long dorsal appendages. Oocytes which had entered the oviduct were discarded along with other stages. These oocytes were observed under a microscope (Axiovert 200M; Zeiss) attached to a spinning disk confocal head (CSU-X1; Yokogawa) controlled by Volocity (PerkinElmer). A Plan-Apochromat objective lens (63x/1.4 numerical aperture) was used with Immersol 518F oil (Zeiss). Images were taken with a Z-slice interval of 0.8 μm and using 40% laser intensity in the red channel, 100% intensity in the green channel. Maximum intensity projections of the Z-stacks are presented as figures and were used for analyses. Images were exported in the TIFF format and signal intensity measurements carried out in ImageJ as followed. The total signal intensities of GFP-Borealin and Rcc1-mChery on the spindle were estimated using the following method. Two areas (S and L) were drawn on the maximum-intensity projection made from Z series of images. Area S includes mainly the central spindle and centromeres or all chromosomes, and Area L includes this area and surrounding region. The total signal intensity over the background was calculated by the formula I_S_ − (N_S_ × [I_L_ − I_S_] / [N_L_ − N_S_]), where I and N are the total pixel intensity and pixel number in the specified area (S or L), respectively.

### Immunostaining and fluorescence *in situ* hybridisation

For immunostaining, ovaries were dissected out of mature flies in methanol as previously described [59]. Sonication was used to disrupt remove the chorion and, which can interfere with antibody penetration. Ovaries were sonicated at 38% amplitude (Vibra Cell VCX500; Sonics) for three 1-second pulses and selected for removal of chorion and vitelline membranes. This was repeated by a further one or two times. The resultant oocytes were washed in 1x PBS, and blocked for 1 hour in blocking buffer (10% foetal calf serum in PBS-T) on a rotator. Oocytes were incubated overnight with primary antibodies diluted in blocking buffer. After washing with PBS-T, fluorophore-conjugated secondary antibodies (Alexa 488, 1:250, or Cy3, 1:1000, in PBS-T) along with DAPI (0.2 µg/ml, Sigma) were added, and incubated for 2 hours in the dark. Finally, oocytes were washed in PBS-T and mounted on microscopy slides in glycerol.

For FISH combined with immunostaining in oocytes, stage 14 oocytes were prepared as for immunostaining, and then post-fixed in 8% formaldehyde as described in [60,61]. An oligonucleotide (CCCGTACTGGT)_4_ for dodecasatellite [40] near centromere 3 was used. 100 pmol of probe was end-labelled with 2 nmol Alexa546-conjugated dUTP (Invitrogen), 16 nmol of unlabelled dTTP (Promega) and 30 units of terminal deoxynucleotidyl transferase (Promega) in 20 μl of transferase buffer at 37°C for one hour The reaction was halted by incubation at 70°C for 10 minutes, and then remaining free dTTP was removed using a G25 Mini Quick spin column (Roche). 4 μl of this labelled oligonucleotide was added to ovaries in 40 μl of hybridisation buffer (0.1 g/ml dextran sulphate, 50% formamide, 3xSSC) at 30°C overnight. After washing twice in washing buffer (50% formamide, 2x SSC, 0.1% Triton X100) at 30°C for total 30 minutes, oocytes were further washed three times in 2xSSC+0.01% Triton X100 and once in PBS. After blocking in PBS+0.1% Triton containing 10% fetal calf serum, the immunostaining procedure was followed for antibody staining.

These slides were then observed via a confocal microscopy (LSM800 on AxioObserver Z1; Zeiss) using a Plan-Apochromat objective lens (63x/1.4 numerical aperture) with Immersol 518F oil (Zeiss). Images spanning an entire spindle were taken with a Z-slice interval of 0.5 μm and Zoom 2.0 (pixel size 0.1 μm). Maximum intensity projections of the Z-stacks are presented as figures.

Expression of GFP-Borr in ovaries were assessed by western blot using a rabbit anti-GFP antibody (1:250, Invitrogen, A11122) and IRDye 800CW conjugated goat anti-mouse IgG antibody (LI-COR). The signals were detected with an Odyssey CLx imaging scanner (LI-COR). MemCode reversible staining kit (Thermo-Fisher) was used to visualise any proteins. The brightness and contrast were adjusted uniformly across the entire area in a linear manner without removing or altering features.

## Acknowledgements

We are grateful to the members of the Ohkura laboratory for their help and for their contributions and for generating reagents, and especially to Emma Peat for her technical assistance. The Bloomington Drosophila Stock Center/Resource Center (National Institutes of Health grants P40OD018537 and 2P40OD010949-10A1) and the Transgenic RNAi Project at Harvard Medical School (National Institutes of Health/National Institute of General Medical Sciences grant R01-GM084947) provided fly stocks and reagents. This work is supported by the Wellcome Trust (098030, 206315, 092076, and 203149) and Biotechnology and Biological Sciences Research Council (BBSRC; BB/S013059). C. Repton received a PhD studentship from BBSRC EastBio Doctoral Training Partnership. The authors declare no competing financial interests.

## Author contributions

CR has contributed to conceptualisation, formal analysis, investigation, analysis and writing the manuscript. CFC and MFAC have contributed to investigation. CS has contributed to investigation and analysis. JR has contributed to supervision and funding acquisition. HO has contributed to supervision, investigation, analysis, funding acquisition and writing the manuscript.

## Supporting information

**S1 Fig.**
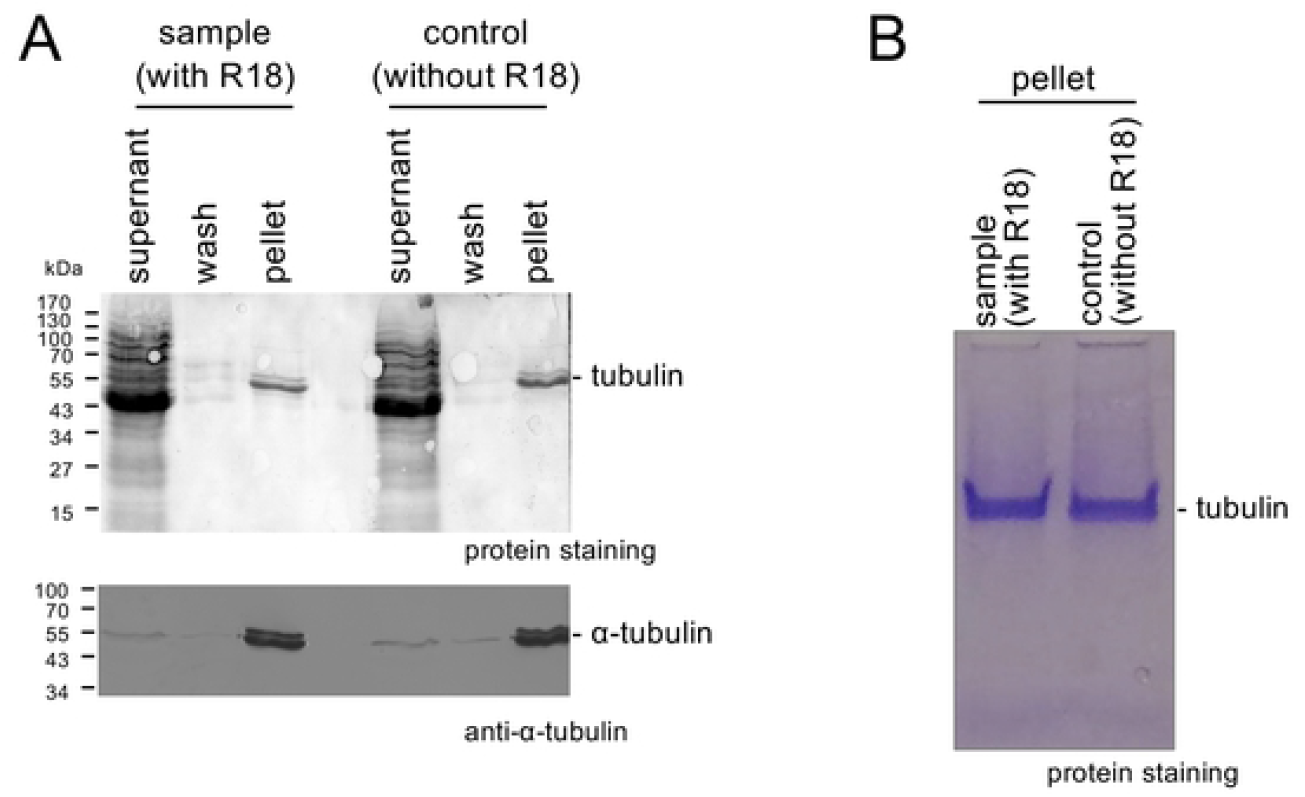
Microtubule co-sedimentation from *Drosophila* ovaries. (A) Microtubules and associated proteins were co-sedimented with or without the 14-3-3 inhibitor R18 from soluble extract of *Drosophila* ovaries. The original supernatant, wash of the original pellet, and the final pellet used for mass-spectrometry were analysed by western blot using an α-tubulin antibody and total protein staining. (B) The final pellets were run on SDS-PAGE and stained with Coomassie for mass-spectrometry. Nearly all tubulin in the extract was found in the pellet fraction, which predominantly consists of tubulin with minimal amounts of other proteins, regardless of the presence or absence of R18.

**S2 Fig.**
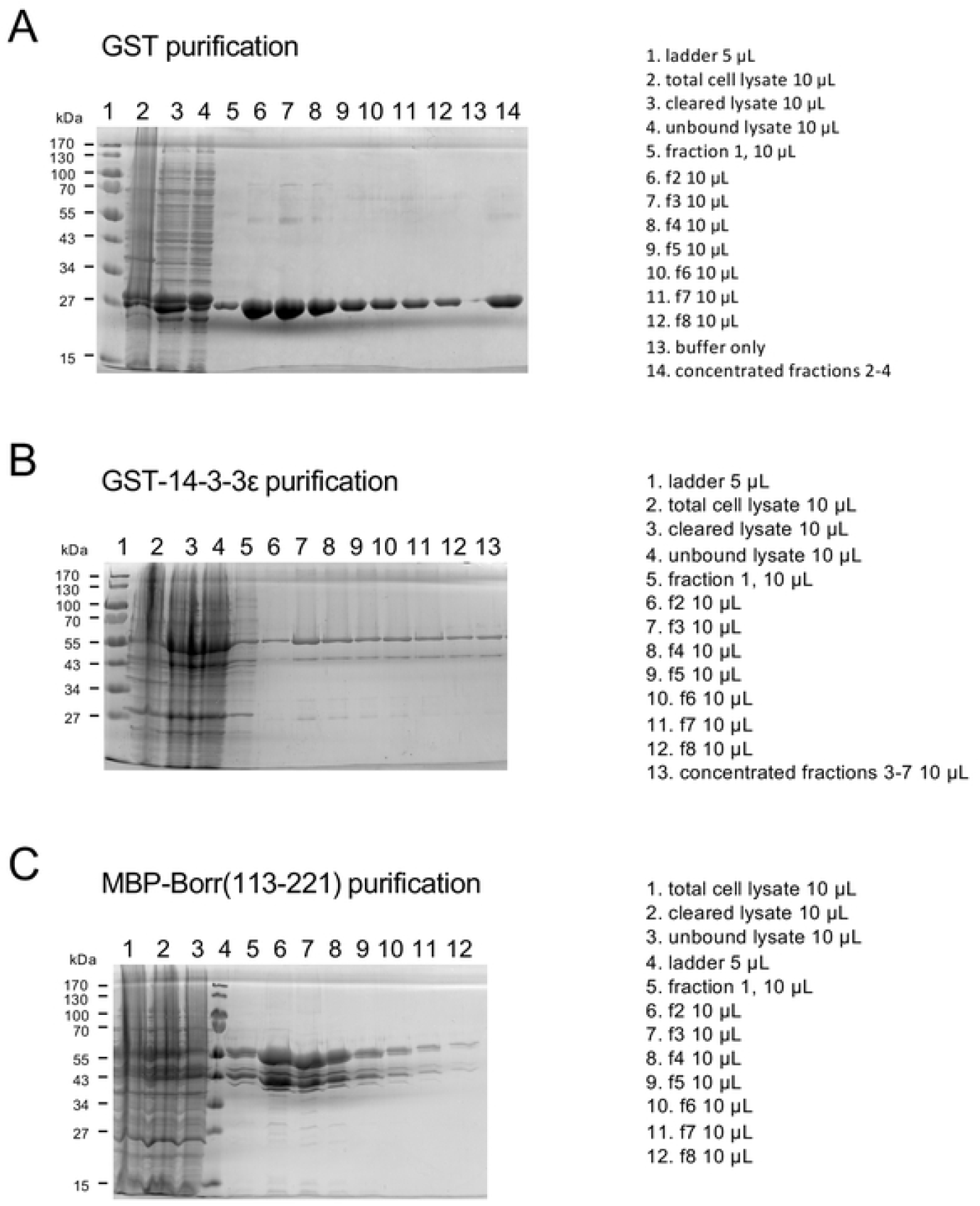
Protein purification of GST, GST-14-3-3ε and MBP-Borr(113-221) from *E. coli*. GST (A), GST-14-3-3ε (B) and MBP-Borr(113-221) (C) were affinity-purified using columns containing S-glutathione beads or amylose beads and eluted in buffer containing glutathione or maltose, respectively, as described in Materials and Methods. Purification intermediates were analysed by SDS-PAGE and stained with Coomassie.

**S1 Table. Mass spectrometry intensity data for all proteins in microtubule fractions with and without R18**

**S2 Table. Raw mass spectrometry intensity data for replicate 1**

**S3 Table. Raw mass spectrometry intensity data for replicate 2**

**S4 Table. Raw mass spectrometry intensity data for replicate 5**

**S5 Table. Raw mass spectrometry intensity data for replicate 6**

**S6 Table. Raw mass spectrometry intensity data for replicate 7**

**S7 Table. Raw mass spectrometry intensity data for replicate 8**

**S8 Table. Amino-acid composition around predicted 14-3-3 binding sites**

The raw numbers of each amino acid at each position around predicted 14-3-3 binding sites, which are used to make Fig. 2C. Position 7 in the table is the phosphorylation site essential for 14-3-3 binding, which corresponds to position 0 in the Fig. 2C. Group 2, 3 and 4 correspond the top 20-40%, 40-60% and 60-80%, respectively.

**S9 Table. Quantification of GFP-Borr signal** Quantification of GFP-Borealin signal, on which Fig 4B is based on.

**S10 Table. Frequencies of bi-oriented homologous centromeres of chromosome 3** The numbers of oocytes showing bi-oriented or mono-oriented homologous centromeres of chromosome 3, on which Fig 4E is based on.

**S11 Table. Parameters used for mass-spectrometry**

## References

1. McKim KS, Hawley RS. Chromosomal Control of Meiotic Cell Division. Science (80-). 1995;270: 1595–1601. doi:10.1126/science.270.5242.1595

2. Cavazza T, Vernos I. The RanGTP pathway: From nucleo-cytoplasmic transport to spindle assembly and beyond. Front Cell Dev Biol. 2016;3. doi:10.3389/fcell.2015.00082

3. Holubcová Z, Blayney M, Elder K, Schuh M. Error-prone chromosome-mediated spindle assembly favors chromosome segregation defects in human oocytes. Obstet Gynecol Surv. 2015;70: 572–573. doi:10.1097/OGX.0000000000000240

4. Schuh M, Ellenberg J. Self-Organization of MTOCs Replaces Centrosome Function during Acentrosomal Spindle Assembly in Live Mouse Oocytes. Cell. 2007;130: 484–498. doi:10.1016/j.cell.2007.06.025

5. Dumont J, Petri S, Pellegrin F, Terret ME, Bohnsack MT, Rassinier P, et al. A centriole-and RanGTP-independent spindle assembly pathway in meiosis I of vertebrate oocytes. J Cell Biol. 2007;176: 295–305. doi:10.1083/jcb.200605199

6. Cesario J, Mckim KS. RanGTP is required for meiotic spindle organization and the initiation of embryonic development in Drosophila. J Cell Sci. 2011;124: 3797–3810. doi:10.1242/jcs.084855

7. Sampath SC, Ohi R, Leismann O, Salic A, Pozniakovski A, Funabiki H. The chromosomal passenger complex is required for chromatin-induced microtubule stabilization and spindle assembly. Cell. 2004;118: 187–202. doi:10.1016/j.cell.2004.06.026

8. Kelly AE, Sampath SC, Maniar TA, Woo EM, Chait BT, Funabiki H. Chromosomal Enrichment and Activation of the Aurora B Pathway Are Coupled to Spatially Regulate Spindle Assembly. Dev Cell. 2007;12: 31–43. doi:10.1016/j.devcel.2006.11.001

9. Kleppe R, Martinez A, Døskeland SO, Haavik J. The 14-3-3 proteins in regulation of cellular metabolism. Semin Cell Dev Biol. 2011;22: 713–719. doi:10.1016/j.semcdb.2011.08.008

10. Denison FC, Paul AL, Zupanska AK, Ferl RJ. 14-3-3 Proteins in Plant Physiology. Semin Cell Dev Biol. 2011;22: 720–727. doi:10.1016/j.semcdb.2011.08.006

11. Pennington K, Chan T, Torres M, Andersen J. The dynamic and stress-adaptive signaling hub of 14-3-3: emerging mechanisms of regulation and context-dependent protein–protein interactions. Oncogene. 2018;37: 5587–5604. doi:10.1038/s41388-018-0348-3

12. Johnson C, Crowther S, Stafford MJ, Campbell DG, Toth R, MacKintosh C. Bioinformatic and experimental survey of 14-3-3-binding sites. Biochem J. 2010;427: 69–78. doi:10.1042/BJ20091834

13. Henriksson ML, Francis MS, Peden A, Aili M, Stefansson K, Palmer R, et al. A nonphosphorylated 14-3-3 binding motif on exoenzyme S that is functional in vivo. Eur J Biochem. 2002;269: 4921–4929. doi:10.1046/j.1432-1033.2002.03191.x

14. Brunet A, Kanai F, Stehn J, Xu J, Sarbassova D, Frangioni J V., et al. 14-3-3 Transits To the Nucleus and Participates in Dynamic Nucleocytoplasmic Transport. J Cell Biol. 2002;156: 817–828. doi:10.1083/jcb.200112059

15. Margolis SS, Walsh S, Weiser DC, Yoshida M, Shenolikar S, Kornbluth S. PP1 control of M phase entry exerted through 14-3-3-regulated Cdc25 dephosphorylation. EMBO J. 2003;22: 5734–5745. doi:10.1093/emboj/cdg545

16. Tang Y, Liu S, Li N, Guo W, Shi J, Yu H, et al. 14-3-3Ζ Promotes Hepatocellular Carcinoma Venous Metastasis By Modulating Hypoxia-Inducible Factor-1Α. Oncotarget. 2016;7: 15854–15867. doi:10.18632/oncotarget.7493

17. Yaffe MB, Rittinger K, Volinia S, Caron PR, Aitken A, Leffers H, et al. The Structural Basis for 14-3-3:Phosphopeptide Binding Specificity. Cell. 1997;91: 961–971. Available: http://dx.doi.org/10.1016/S0092-8674(00)80487-0

18. Beaven R, Bastos RN, Spanos C, Romé P, Fiona Cullen C, Rappsilber J, et al. 14-3-3 regulation of Ncd reveals a new mechanism for targeting proteins to the spindle in oocytes. J Cell Biol. 2017;216: 3029–3039. doi:10.1083/jcb.201704120

19. De S, Kline D. Evidence for the requirement of 14-3-3eta (YWHAH) in meiotic spindle assembly during mouse oocyte maturation. BMC Dev Biol. 2013;13. doi:10.1186/1471-213X-13-10

20. Lawrence CJ, Dawe RK, Christie KR, Cleveland DW, Dawson SC, Endow SA, et al. A standardized kinesin nomenclature. J Cell Biol. 2004;167: 19–22. doi:10.1083/jcb.200408113

21. Hatsumi M, Endow SA. Mutants of the microtubule motor protein, nonclaret disjunctional, affect spindle structure and chromosome movement in meiosis and mitosis. J Cell Sci. 1992;101: 547–559.

22. Mountain V, Simerly C, Howard L, Ando A, Schatten G, Compton DA. Cross-links Microtubules in the Mammalian Mitotic Spindle. J Cell Biol. 1999;147: 351–365.

23. Syred HM, Welburn J, Rappsilber J, Ohkura H. Cell cycle regulation of microtubule interactomes: multi-layered regulation is critical for the interphase/mitosis transition. Mol Cell Proteomics. 2013;12: 3135–47. doi:10.1074/mcp.M113.028563

24. Wang B, Yang H, Liu YC, Jelinek T, Zhang L, Ruoslahti E, et al. Isolation of high-affinity peptide antagonists of 14-3-3 proteins by phage display. Biochemistry. 1999;38: 12499–12504. doi:10.1021/bi991353h

25. Eisa AA, D. S, Detwiler A, Gilker E, Ignatious AC, Vijayaraghavan S, et al. YWHA (14-3-3) protein isoforms and their interactions with CDC25B phosphatase in mouse oogenesis and oocyte maturation. BMC Dev Biol. 2019;19: 1–22. doi:10.1186/s12861-019-0200-1

26. Wu C, Muslin AJ. Role of 14-3-3 proteins in early Xenopus development. Mech Dev. 2002;119: 45–54. doi:10.1016/S0925-4773(02)00287-3

27. Szklarczyk D, Gable AL, Nastou KC, Lyon D, Kirsch R, Pyysalo S, et al. The STRING database in 2021: Customizable protein-protein networks, and functional characterization of user-uploaded gene/measurement sets. Nucleic Acids Res. 2021;49: D605–D612. doi:10.1093/nar/gkaa1074

28. Douglas ME, Davies T, Joseph N, Mishima M. Aurora B and 14-3-3 Coordinately Regulate Clustering of Centralspindlin during Cytokinesis. Curr Biol. 2010;20: 927–933. doi:10.1016/j.cub.2010.03.055

29. Hein JB, Hertz EPT, Garvanska DH, Kruse T, Nilsson J. Distinct kinetics of serine and threonine dephosphorylation are essential for mitosis. Nat Cell Biol. 2017;19: 1433–1440. doi:10.1038/ncb3634

30. Colombié N, Cullen CF, Brittle AL, Jang JK, Earnshaw WC, Carmena M, et al. Dual roles of incenp crucial to the assembly of the acentrosomal metaphase spindle in female meiosis. Development. 2008;135: 3239–3246. doi:10.1242/dev.022624

31. Wignall SM, Villeneuve AM. Lateral microtubule bundles promote chromosome alignment during acentrosomal oocyte meiosis. Nat Cell Biol. 2009;11: 839–844. doi:10.1038/ncb1891

32. Carmena M, Wheelock M, Funabiki H, Earnshaw WC. The chromosomal passenger complex (CPC): From easy rider to the godfather of mitosis. Nat Rev Mol Cell Biol. 2012;13: 789–803. doi:10.1038/nrm3474

33. Radford SJ, Jang JK, McKim KS. The Chromosomal Passenger Complex Is Required for Meiotic Acentrosomal Spindle Assembly. Genetics. 2012;192: 417–429. doi:10.1534/genetics.112.143495

34. Romé P, Ohkura H. A novel microtubule nucleation pathway for meiotic spindle assembly in oocytes. J Cell Biol. 2018;217: 3431–3445. doi:10.1083/jcb.201803172

35. Wang L, Defosse T, Jang JK, Battaglia RA, Wagner VF, Mckim KS. Borealin directs recruitment of the CPC to oocyte chromosomes and movement to the microtubules. J Cell Biol. 2021;220: e202006018. doi:doi.org/10.1083/jcb.2020060181of

36. Costa MFA, Ohkura H. The molecular architecture of the meiotic spindle is remodeled during metaphase arrest in oocytes. J Cell Biol. 2019;218: 2854–2864. doi:10.1083/jcb.201902110

37. Jeyaprakash AA, Klein UR, Lindner D, Ebert J, Nigg EA, Conti E. Structure of a Survivin-Borealin-INCENP Core Complex Reveals How Chromosomal Passengers Travel Together. Cell. 2007;131: 271–285. doi:10.1016/j.cell.2007.07.045

38. Bekier ME, Mazur T, Rashid MS, Taylor WR. Borealin dimerization mediates optimal CPC checkpoint function by enhancing localization to centromeres and kinetochores. Nat Commun. 2015;6. doi:10.1038/ncomms7775

39. Resnick TD, Satinover DL, MacIsaac F, Stukenberg PT, Earnshaw WC, Orr-Weaver TL, et al. INCENP and Aurora B Promote Meiotic Sister Chromatid Cohesion through Localization of the Shugoshin MEI-S332 in Drosophila. Dev Cell. 2006;11: 57–68. doi:10.1016/j.devcel.2006.04.021

40. Abad JP, Carmena M, Baars S, Saunders RDC, Glover DM, Ludeña P, et al. Dodeca satellite: A conserved G+C-rich satellite from the centromeric heterochromatin of Drosophila melanogaster. Proc Natl Acad Sci U S A. 1992;89: 4663–4667. doi:10.1073/pnas.89.10.4663

41. Radford SJ, Go AMM, Mckim KS. Spindle Symmetry and Chromosome Organization. Genetics. 2017;205: 517–527. doi:10.1534/genetics.116.194647

42. Trivedi P, Zaytsev A V., Godzi M, Ataullakhanov FI, Grishchuk EL, Stukenberg PT. The binding of Borealin to microtubules underlies a tension independent kinetochore-microtubule error correction pathway. Nat Commun. 2019;10. doi:10.1038/s41467-019-08418-4

43. Ashburner M. Drosophila: A Laboratory Handbook and Manual. Cold Spring Harbour Laboratory Press; 1989.

44. Venken KJT, He Y, Hoskins RA, Bellen HJ. P[acman]: A BAC transgenic platform for targeted insertion of large DNA fragments in D. melanogaster. Science (80-). 2006;314: 1747–1751. doi:10.1126/science.1134426

45. Perkins LA, Holderbaum L, Tao R, Hu Y, Sopko R, McCall K, et al. The transgenic RNAi project at Harvard medical school: Resources and validation. Genetics. 2015;201: 843–852. doi:10.1534/genetics.115.180208

46. Hughes JR, Meireles AM, Fisher KH, Garcia A, Antrobus PR. A Microtubule Interactome : Complexes with Roles in Cell Cycle and Mitosis. PLoS Biol. 2008;6: 0785–0795. doi:10.1371/journal.pbio.0060098

47. Shevchenko A, Tomas H, Havlis J, Olsen J V, Mann M. In-gel digestion for mass spectrometric characterization of proteins and proteomes. Nat Protoc. 2007;1: 2856–2860. doi:10.1038/nprot.2006.468

48. Rappsilber J, Ishihama Y, Mann M. Stop and Go Extraction Tips for Matrix-Assisted Laser Desorption /Ionization, Nanoelectrospray, and LC /MS Sample Pretreatment in Proteomics. Anal Chem. 2003;75: 663–670.

49. Olsen J V, Macek B, Lange O, Makarov A, Horning S, Mann M. Higher-energy C-trap dissociation for peptide modification analysis. Nat Methods. 2007;4: 709–712. doi:10.1038/NMETH1060

50. Cox J, Mann M. MaxQuant enables high peptide identification rates, individualized p.p.b.-range mass accuracies and proteome-wide protein quantification. Nat Biotechnol. 2008;26: 1367–1372. doi:10.1038/nbt.1511

51. Cox J, Neuhauser N, Michalski A, Scheltema RA, Olsen J V, Mann M. Andromeda : A Peptide Search Engine Integrated into the MaxQuant Environment. J Proteome Res. 2011;10: 1794–1805.

52. Vizcaíno JA, Côté RG, Csordas A, Dianes JA, Fabregat A, Foster JM, et al. The Proteomics Identifications (PRIDE) database and associated tools: Status in 2013. Nucleic Acids Res. 2013;41: 1063–1069. doi:10.1093/nar/gks1262

53. St. Pierre SE, Ponting L, Stefancsik R, Mcquilton P, Consortium TF. FlyBase 102 — advanced approaches to interrogating FlyBase. Nucleic Acids Res. 2014;42: 780–788. doi:10.1093/nar/gkt1092

54. Team RC. R: A language and environment for statistical computing. Vienna, Austria.: R Foundation for Statistical Computing; 2017. Available: https://www.r-project.org/.

55. Wickham H. ggplot2: Elegant Graphics for Data Analysis. Springer-Verlag New York.; 2009.

56. Madeira F, Tinti M, Murugesan G, Berrett E, Stafford M, Toth R, et al. Sequence analysis 14-3-3-Pred : improved methods to predict 14-3-3-binding phosphopeptides. Bioinformatics. 2015;31: 2276–2283. doi:10.1093/bioinformatics/btv133

57. Xue Y, Liu Z, Cao J, Ma Q, Gao X, Wang Q, et al. GPS 2.1: Enhanced prediction of kinase-specific phosphorylation sites with an algorithm of motif length selection. Protein Eng Des Sel. 2011;24: 255–260. doi:10.1093/protein/gzq094

58. Crooks G, Hon G, Chandonia J, Brenner S. NCBI GenBank FTP Site\nWebLogo: a sequence logo generator. Genome Res. 2004;14: 1188–1190. doi:10.1101/gr.849004.1

59. Cullen CF, Ohkura H. Msps protein is localized to acentrosomal poles to ensure bipolarity of Drosophila meiotic spindles. Nat Cell Biol. 2001;3: 637–642.

60. Meireles AM, Fisher KH, Colombie N, Wakefield JG, Ohkura H. Wac: a new Augmin subunit required for chromosome alignment but not for acentrosomal microtubule assembly in female meiosis. J Cell Biol. 2009;184: 777–784. doi:10.1083/jcb.200811102

61. Nikalayevich E, Ohkura H. The NuRD nucleosome remodelling complex and NHK-1 kinase are required for chromosome condensation in oocytes. J Cell Sci. 2015;128: 566–575. doi:10.1242/jcs.158477

